# A practical tool for Maximal Information Coefficient analysis

**DOI:** 10.1101/215855

**Authors:** Davide Albanese, Samantha Riccadonna, Claudio Donati, Pietro Franceschi

## Abstract

**Background:** The ability of finding complex associations in large omics datasets, assessing their significance, and prioritizing them according to their strength can be of great help in the data exploration phase. Mutual Information based measures of association are particularly promising, in particular after the recent introduction of the TIC_e_ and MIC_e_ estimators, which combine computational efficiency with good bias/variance properties. Despite that, a complete software implementation of these two measures and of a statistical procedure to test the significance of each association is still missing.

**Findings:** In this paper we present MICtools, a comprehensive and effective pipeline which combines TIC_e_ and MIC_e_ into a multi-step procedure that allows the identification of relationships of various degrees of complexity. MICtools calculates their strength assessing statistical significance using a permutation-based strategy. The performances of the proposed approach are assessed by an extensive investigation in synthetic datasets and an example of a potential application on a metagenomic dataset is also illustrated.

**Conclusions:** We show that MICtools, combining TIC_e_ and MIC_e_, is able to highlight associations that would not be captured by conventional strategies. MICtools is implemented in Python, and is available for download at https://github.com/minepy/mictools.

## Introduction

With the growing popularity of high throughput quantitative technologies it is now common to characterize living systems by measuring thousands of variables over a wide range of conditions. In these large datasets, the number of potential associations between variables is enormous. Computational and statistical methods should be able to highlight the significant ones (striking a balance between flexibility and statistical robustness), and to prioritize the more relevant for downstream analysis. Traditionally, the presence of a potential relationship between two variables *X* and *Y* is assessed on the basis of a certain measure of association, that is often able to reveal specific types of relationships, but is blind to others. Then, once the measure is computed, its significance is tested against the null hypothesis of no association. For linear associations, the Pearson correlation coefficient is the natural choice, while the Spearman’s rank coefficient represents a more flexible alternative for general monotonic relationships. In the exploratory analysis of datasets produced by modern-omics technologies this conventional approach show its limits, because a huge number of potential associations needs to be screened without any *a priori* information on their form. In these cases, it would be desirable to use a measure of dependence that ranks the relationships according to their strength, regardless of the type of association. A measure with this property has been defined *equitable* [1] and a consistent mathematical framework for the definition of equitability has been proposed [2, 3, 4, 5, 6]. The second challenge faced in the unsupervised screening of large datasets is that the number of associations to be tested is usually huge and the statistical assessment of significance has to face well known multiplicity issues [7, 8].

Recently, a family of measures based on the concept of mutual information has been proposed, and one of the most popular member of this family, the Maximal Information Coefficient (MIC), has been shown to satisfy the equitability requirement [1]. Unfortunately, MIC suffers of lack of power [9], and its heuristic estimator, APPROX-MIC, is computationally demanding [5]. These two drawbacks have severely hampered the application of MIC to large datasets. In order to overcome these limitations, two new MIC-based measures, the MIC_e_— a consistent estimator of the MIC population value (MIC_*_) — and the related TIC_e_ (total information coefficient) statistics have been proposed [5]. Both quantities can be calculated more efficiently than APPROX-MIC and have better bias/variance properties [5]. In particular, TIC_e_ is characterized by high power, which has been obtained at the expenses of equitability, while MIC_e_ performs better on this side, showing reduced performances in terms of power. These two MIC-based measures, then, compensate each other and their combination is extremely promising as data exploration tool. In particular, a two step procedure can be applied, where TIC_e_ is used to perform efficiently a high throughput screening of all the possible pairwise relationships and assess their significance, while MIC_e_ is used to rank the subset of significant associations in terms of strength [5]. Despite the potential of this approach, an efficient software implementation of these two measures and of a statistical procedure to test the significance of each association controlling multiplicity issues is still lacking.

Here we present MICtools, an open-source and easy-to-use software providing:

- an efficient implementation of TIC_e_ and MIC_e_ estimators[10];
- a permutation-based strategy for estimating TIC_e_ empirical *p* values;
- several methods for multiple testing correction, including the Storey’s *q* value to control the false discovery rate (FDR);
- the MIC_e_ estimates for each association called significant.

## Methods

MICtools implements a multi-step procedure to identify relevant associations amongst a large number of variables, assess their statistical significance and rank them according to the strength of the relationship. Starting from *M* variable pairs *x*_*i*_ and *y*_*i*_ measured in *n* samples, the procedure can be broken into 4 steps (Figure 1):

i. estimating the empirical TIC_e_ null distribution by permutations;
ii. computing TIC_e_ statistics and its empirical *p* values for each variable pairs;
iii. applying a multiple testing correction strategy in order to control the family-wise error rate (FWER) or the FDR [11];
iv. using MICe to estimate the strength of the relationships called significant.

**Figure 1.**
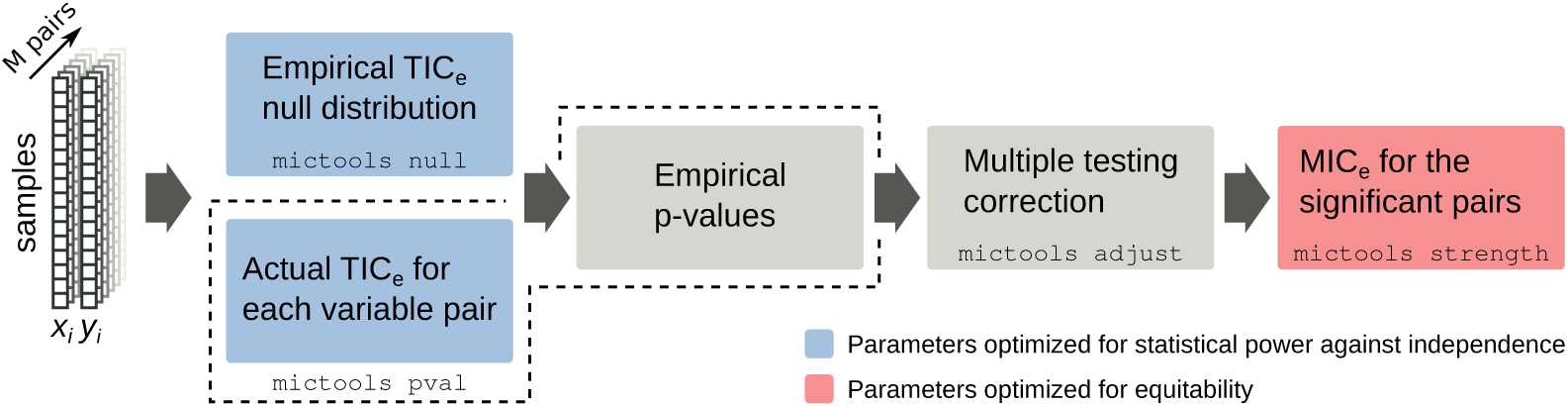
The MICtools pipeline. Each step is implemented as a subcommand of the `mictools` main command. `mictools` null estimates the empirical TIC_e_ null distribution of the *M* variable pairs (*x*_*i*_,*y*_*i*_). `mictools` pval computes the TIC_e_ values and estimates their *p* values (boxes withing the dashed line). The multiple testing correction is performed by `mictools` adjust. Finally, `mictools` strength estimates the MIC_e_ value for the subset of significant relationships. The color of the boxes highlights the criterion used for parameter optimization.

The pipeline can be run as a sequence of subcommands implemented into the main command `mictools` (Figure 1).

### The empirical TIC_e_ null distribution

Since TIC_e_ depends only on the rank-order of the vectors *x*_*i*_ and *y*_*i*_ [1], the empirical null distribution can be estimated, for a given sample size and set of parameters, by performing *R* permutations of the elements of the vectors *y*_*i*_ and by calculating the set of null TIC_e_ statistics 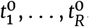. Two parameters control the estimation of the null distribution of TICe: the parameter *B* controlling the maximal-allowed grid resolution and the number of permutations *R*. In the current implementation, *B* was set to the default value 9, which guarantees good performances in terms of statistical power against independence in most situations [12]. However, different values of *B* can be chosen: for example, *B* = 4 for less complex alternative hypothesis, *B* = 12 for more complex associations [12]. With regards to the number of permutations, instead, the results obtained on the synthetic datasets (see Additional File 2, Figures A2 and A3 and Additional File 1, Table A2) empirically indicate that 200,000 permutations represent a reasonable choice in most scenarios.

### Computing the TIC_e_ and its associated empirical *p* values for each variable pair

The total information coefficient is computed for each (non permuted) variable pair, obtaining a set of TIC_e_ values *t*_*i*_ (with *i* = {1,…,*M*}). For each *t*_*i*_, the *p* value *p*_*i*_ is estimated as the fraction of values of the empirical null distribution that exceeds *t*_*i*_ [13]:

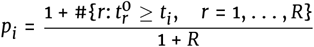

### Multiple testing correction

Considering the large number of tests of independence performed, it is necessary to correct the *p* values for multiplicity. This can be done either by controlling the FWER or the FDR. MICtools implements several state-of-the-art strategies to accomplish this task. For all the examples presented here we have used the Storey’s method for estimating the *q* values to control the FDR [7]. Briefly, setting a *q*-value cut-off to 0.05, we accept a FDR of at most 5%.

### Computing the MICe on the significant relationships

Finally, the strength of the associations that pass the significance threshold is estimated using the MIC_e_ estimator. In this case, we define the the *B* parameter as a function of the number of samples *n, B(n)* = *n*^∝^ [1]. The default values are optimized for equitability [6] and summarized in Table 1.

**Table 1.** The default values of the α parameter vary according to the number of samples.

| Number of samples $n$ | $\alpha$ parameter |
| --- | --- |
| $n < 25$ | 0.85 |
| $25 \leq n < 50$ | 0.80 |
| $50 \leq n < 250$ | 0.75 |
| $250 \leq n < 500$ | 0.70 |
| $500 \leq n < 1000$ | 0.65 |
| $1,000 \leq n < 2,500$ | 0.60 |
| $2,500 \leq n < 5,000$ | 0.55 |
| $5,000 \leq n < 10,000$ | 0.50 |
| $10,000 \leq n < 40,000$ | 0.45 |
| $n > 40,000$ | 0.40 |

## Findings

Two synthetic datasets (SD1 and SD2) were created in order to assess (i) the statistical power (or recall, i.e. the fraction of non-independent relationships that were recovered at a given significance level) and (ii) the ability to control the FDR. The analyses were performed varying the number of samples (SD1) and the effect chance [14], i.e. the percentage of non independent variable pairs (SD2). Both datasets contain a set of independent variables and a fixed number of variable pairs *X* and *Y* related by associations in the form *Y* = *f(X)* + *η*, where *f*(*X*) is a function and *η* is a noise term. To characterize the performances of MICtools in presence of associations that could not be described by a function, a series of Madelon datasets [15,16] was also analyzed. Finally, the proposed pipeline was applied to the analysis of an environmental/metagenomic dataset which has been recently made available within the Tara project, a global-scale characterization of plankton using high throughput metagenomic sequencing [17].

### Synthetic datasets

The SD1 dataset contains 60,000 associations between variable pairs *X* and *Y*. The effect chance was set to 1%. The relationships between the 600 non-independent variable pairs were randomly chosen among 6 different types of functional associations, namely cubic, exponential (2^x^), line, parabola, sigmoid, and spike (see Table S3 in [1]). The noise term *η* is a random variable with uniform distribution in the range of f(X) multiplied by an intensity factor *k*_*η*_. Different values of *k*_*η*_ were chosen randomly among 18,000 values obtained joining the following three sequences: the first ranging from 0.05 to 1 (with steps of 0.0001), the second ranging from 1 to 2 (with steps of 0.0002), and the third ranging from 2 to 9 (with steps of 0.002). Using these values, the coefficients of determination (*R*^2^) between *Y* and the noiseless function *f(X)* ranges approximately from 0 to 1. The remaining 99% (59,400) associations were defined with *X* and *Y* randomly generated from a uniform distribution between 0 and 1. To characterize the effect of the sample size, we created 20 replicates of SD1 for an increasing number of samples (*n* ∈ {25, 50, 100, 250, 1,000}), for a total of 100 datasets. Considering that the fraction of true positive associations was known, this design of experiment allowed us to quantify the statistical power and the performances in terms of FDR of the proposed pipeline. The results for 2 × 10^5^ permutations are summarized in Figure 2 and in the Additional File 1, Table A1. The dependence of the power and of the number of false positives (FP) from the number of samples are shown in Figure 2a and 2b. The power increases with the number of samples reaching 75% for a sample size of 100. As expected, considering that we used the Storey’s *q* value as a strategy to control the FDR, also the number of false positive grows for increasing sample size (Figure 2b) to keep the false discovery rate constant (0.05 in this case). Figure 2c shows the observed FDR, which is almost equal to the expected value of 0.05 for all samples sizes. In Figure 2d we show the values of MIC_e_ as a function of the coefficient of determination (R^2^) between *Y* and the noiseless function *f*(*X*) for the associations that pass the significance filter (i.e. associations with *q* values <0.05). As expected, MIC_e_ and R^2^ were always linearly correlated, especially for the larger sample sizes [5] (Figure 2d, upper panel). Moreover, we found that, for small sample sizes, only relationships with relatively high values of *R*^2^ passed the significance filter. This effect decreases with increasing number of samples, showing that the pipeline is able to identify relationships with more noise, provided that a sufficient number of experimental points is available. This effect is clearly visible in Figure 2e, where we show the statistical power as a function of the strength of the relationships for different sample sizes. While on less noisy associations (having *R*^2^ close to 1) the pipeline shows high power also for smaller sample sizes, a high number of samples is needed to attain high power for very noisy relationships (having *R*^2^ close to 0). Upon closer inspection, the panel d in Figure 2 also shows that the power depends on the form of the association. For instance, red points (corresponding to cubic functional forms) are hardly visible for sample sizes smaller than 100, while sigmoidal, linear and exponential relationships can be identified for all sample sizes, albeit with a power that depends on the amount of noise. This finding can be easily interpreted considering that more complex relationships (e.g. polynomials of higher order) are defined by a higher number of parameters that makes them more difficult to distinguish from random associations if the number of points is limited. A more clear representation of this phenomenon is included in Additional File 2 (Figure A1). Moreover, the downward bias in terms of equitability, especially for the more complex relationships (Figure 2d and A1) is a result of the core approximation algorithm EQUICHARCLUMP, which speeds up the computation of MIC_e_ [18, 5]. The EQUICHARCLUMP parameter *c* controls the coarseness of the discretization in the grid search phase and by default it is set to 5, providing good performance in most settings [12].

**Figure 2.**
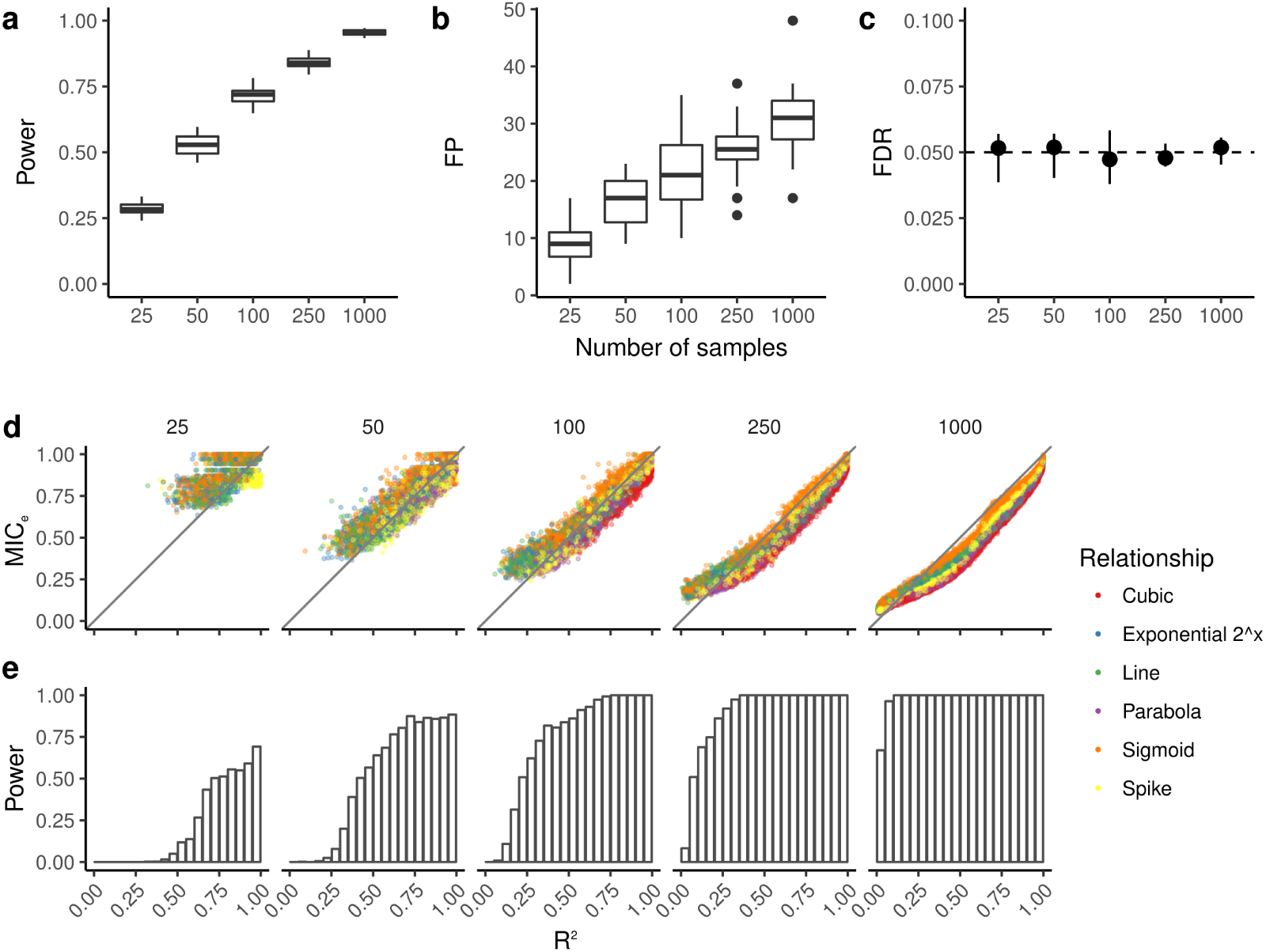
Overview of the CaDRReS framework. Analysis on SD1 dataset at the 0.05 significance level. (a) Statistical power, (b) number of false positives (FP), and (c) false discovery rate (FDR) at varying number of samples *n*. Each range represents the results of the 20 replicates. (d) MIC_e_ values and (e) statistical power at different levels of *R*^2^, for increasing number of samples (from 25 to 1,000, plots from left to right). Only significant relationships, i.e. relationships with *q* < 0.05, are shown.

As already anticipated, SD1 was also used to investigate the dependence of the performances of MICtools on the number of independent permutations used to estimate the empirical null distribution. Figure A2 and A3 (Additional File 2) shows the FDR and the power as a function of the number of samples and of the number of permutations. The plots indicate that for all the combinations of the two parameters the measured FDR was consistent with the expected value 0.05 (Additional File 2, Figure A2 and Additional File 1, Table A2) and that the true value is always included in the shaded interquartile area. As expected, the variability is stronger for the smaller dataset (25 samples), but also with such a small number of samples above 2 × 10^5^ permutations the median measured FDR stabilizes around 0.05. Figure A3a in the Additional File 2 shows the expected increase of power with the number of samples, from 0.25 to almost 1. The median values does not show a strong dependence on the number of permutations. Figure A3b indicates that below 100 samples at least 2 × 10^5^ permutations are needed to obtain stable values of power, and that its variability is anyway larger for small sample sets. All these evidences, however, support the choice of 200,000 as a default value for the number of permutations.

The dataset SD2 was generated to characterize how the effect chance, i.e. the fraction of non-random associations, affected the performances of MICtools. Similarly to dataset SD1, SD2 contains a subset of variable pairs *X* and *Y* related by associations of the form *Y* = *f(X)* + *η*, where *η* was defined as in SD1. The number of samples was fixed to *n* = 100 and the total number of associations was 60,000. For each effect chance value (1%, 2%, 5%, 10%, 20% and 50%) we generated 20 independent datasets, for a total of 120. The power, number of False Positives (FP) and FDR as a function of the effect chance are shown in Figure 3, panels a, b and c, respectively (see also Additional File 1, Table A3). In Figure 3a, we can observe that the statistical power grows with the effect chance, while the actual FDR remains constant. In fact, an increase of effect chance corresponds to a decrease of the fraction of relationships for which the null is true, π_0_ (effect chance = 1 - π_0_). Consequently, an increase of the *p*-value threshold and therefore a growth of power is expected in order to maintain the FDR cutoff constant [7, 14].

**Figure 3.**
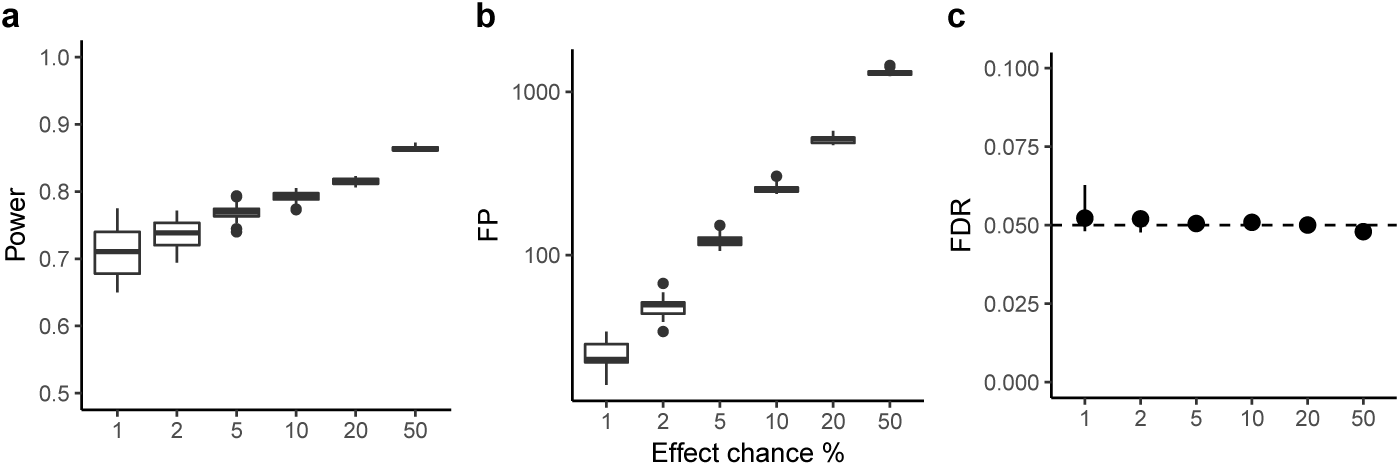
Analysis on SD_2_ dataset at the 0.05 significance level. (a) Statistical power, (b) number of False positive (FP), and (c) False discovery rate (FDR) for increasing effect chance. Each range represents 20 replicated datasets.

## The Madelon classification dataset

The analysis of SD1 and SD2 datasets demonstrates that MICtools is able to identify the relationships described by analytic functions with additive noise. However, more general forms of non-random associations are possible. Consider, for instance, the presence of clusters that might indicate the presence of subpopulations. To test the ability of MICtools to identify this type of associations, we created 7 datasets with an increasing number of samples *n* ∈ {50, 250, 500, 1,000, 2,500, 5,000} with a structure similar to the Madelon binary classification dataset [16, 15] (http://archive.ics.uci.edu/ml/machine-learning-databases/madelon/Dataset.pdf) using the `datasets.make_ciassification()` function available in the scikit-learn library [19]. Each dataset contains 4 clusters (two for each class), placed on the vertices of a five dimensional four-sided hypercube. Each cluster was composed by normally distributed points (σ = 1). The five dimensions defining the hypercube constitutes the 5 “informative” features. Other 15 “redundant” features were generated as random linear combinations of the informative features and added to the dataset. Finally, 180 random variables without predictive power were added, for a total of 200. In this type of setting, the number of associations to be tested was 19900 = (200 × 199)/2. Among them, 190 are “real” (the relationships between the variables belonging to the “informative” and “redundant”). Figure 4a summarizes the results of the analysis. Panel (a) shows the association called significant (*q*-value cutoff set to 0.05) on a Hive plot [20] as a functions of the number of samples. Each branch of the Hives represents a type of variable (informative: 5 variables; redundant: 15; random: 180), the blue lines identify true positives (associations between non-independent variables correctly identified), while false positives (incorrectly identified associations between independent variables) are marked in red. This representation clearly shows that, as expected, the number of true positives increases with the number of samples. A more quantitative representation of the effect of the number of samples on the number of false negatives (non-independent associations incorrectly rejected) is shown in panel (b). Again, an increase in the number of samples is beneficial, because it reduces the number of false negatives. The last panel (c) of Figure 4 shows the effect of n on the FDR, which is always approximately constant and very close to the theoretical value of 0.05.

**Figure 4.**
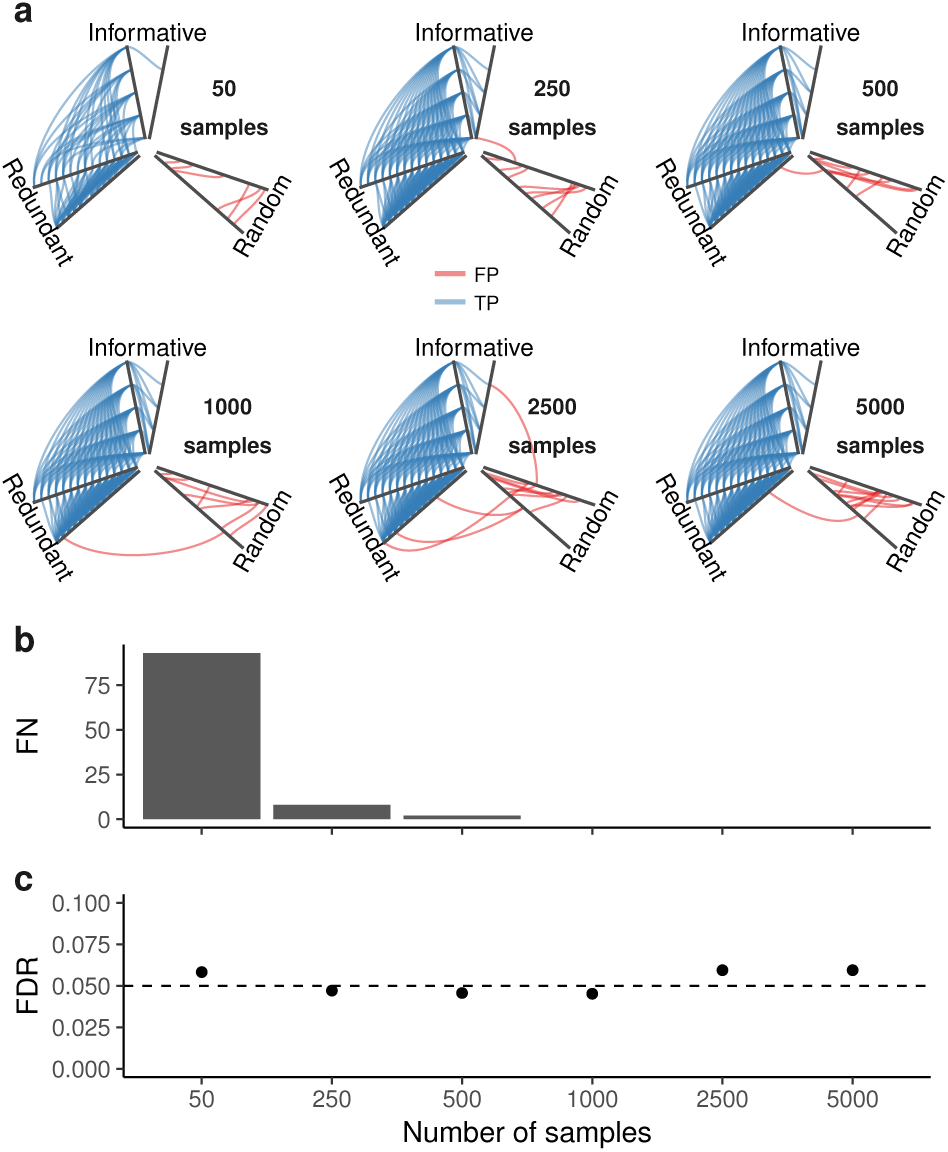
Madelon dataset. (a) Hive plots of the detected association for increasing number of samples. The variables are grouped as “informative” (5), “redundant” (15), and “random” (180). True positives (associations between non-independent variables passing the significance test) are in blue; false positives (associations between independent variable passing the significance test) in red. (b) Number of false negatives (FN), i.e. number of non-independent variable pairs which were not detected as significant, and (c) false discovery rate (FDR) as a function of the number of samples.

On the bases of these results, we conclude that also with a relatively low number of samples MICtools is able to identify in an efficient way non functional associations typical of cluster structures. It is interesting to note that the associations among the informative variables started to be recovered when at least 250 samples were considered, while the associations between informative/redundant and redundant/redundant variables were significant also for lower number of samples (50). This apparently odd behavior is due to the different nature of the association among the variables. Binary associations among informative variables are indeed characterized by the presence of clusters, while redundant associations are constructed by linear combinations. In accordance to the results discussed for SD1, the statistical power of the procedure depends on the type of association and with a lower number of samples the results are biased towards less complex association patterns.

## Identifying ecological niches: the Tara dataset

The Tara Oceans project is a large multinational effort for the study of plankton at a global scale [17]. Within the project, a large study of the microbiota of water samples from the oceans characterized using metagenomic techniques has been recently made available. To illustrate the added value of using MICtools to analyze such large datasets, we downloaded the annotated 16S _mi_tags [21] OTU count table of 139 water samples from http://ocean-microbiome.embl.de/companion.html, together with the accompanying metadata on temperature and chemical composition [22]. MICtools was used to identify the existence of significant relationships between the environmental variables and the taxonomic composition of the microbiota. The genus relative abundances, the environment variables and the samples metadata are available in the Additional File 1, tables A4, A5 and A6 respectively. By using a *q*-value cutoff of 0.01 we found significant associations between the relative abundances of 279 taxa with water temperature and of 287 taxa with oxygen (Figure 5, panels b and c, respectively). To highlight the novel information provided by MICtools, Spearman’s rank correlation coefficients and their associated p values were also calculated as in [23] (the default for the `cor.test()` function in the R environment). By using the Spearman’s coefficient alone we could identify a subset of the relations identified by MICtools, namey 194 taxa were associated with temperature and 191 taxa were associated with oxygen concentration, respectively. Conversely, almost all relationships identified with Spearman’s correlation were also identified by MICtools. While the Spearman’s coefficient based approach identified associations well described by monotonic functions (Figure 5e and 5f), MICtools was able to highlight the presence of more complex relationships between the taxa and the environmental parameters. As an example, we found a sharp increase of the Alcaligenaceae genus at oxygen concentration of 200 μmol kg^-1^ (Figure 5d) and a slow increase of the Sphingomonadaceae genus as a function of the temperature. In both cases, highlighting the samples on the bases of their specific aquatic layer of reference it is possible to see that the complex aggregation patterns identified by MICtools are associated to specific ecological niches. These results show the advantage of the use of the proposed approach as an automatic screening tool in the data exploration phase. The lists of the relationships identified by MICtools and by the Spearman-based procedure are available in the Additional File 1, Tables A7 and A8, respectively.

**Figure 5.**
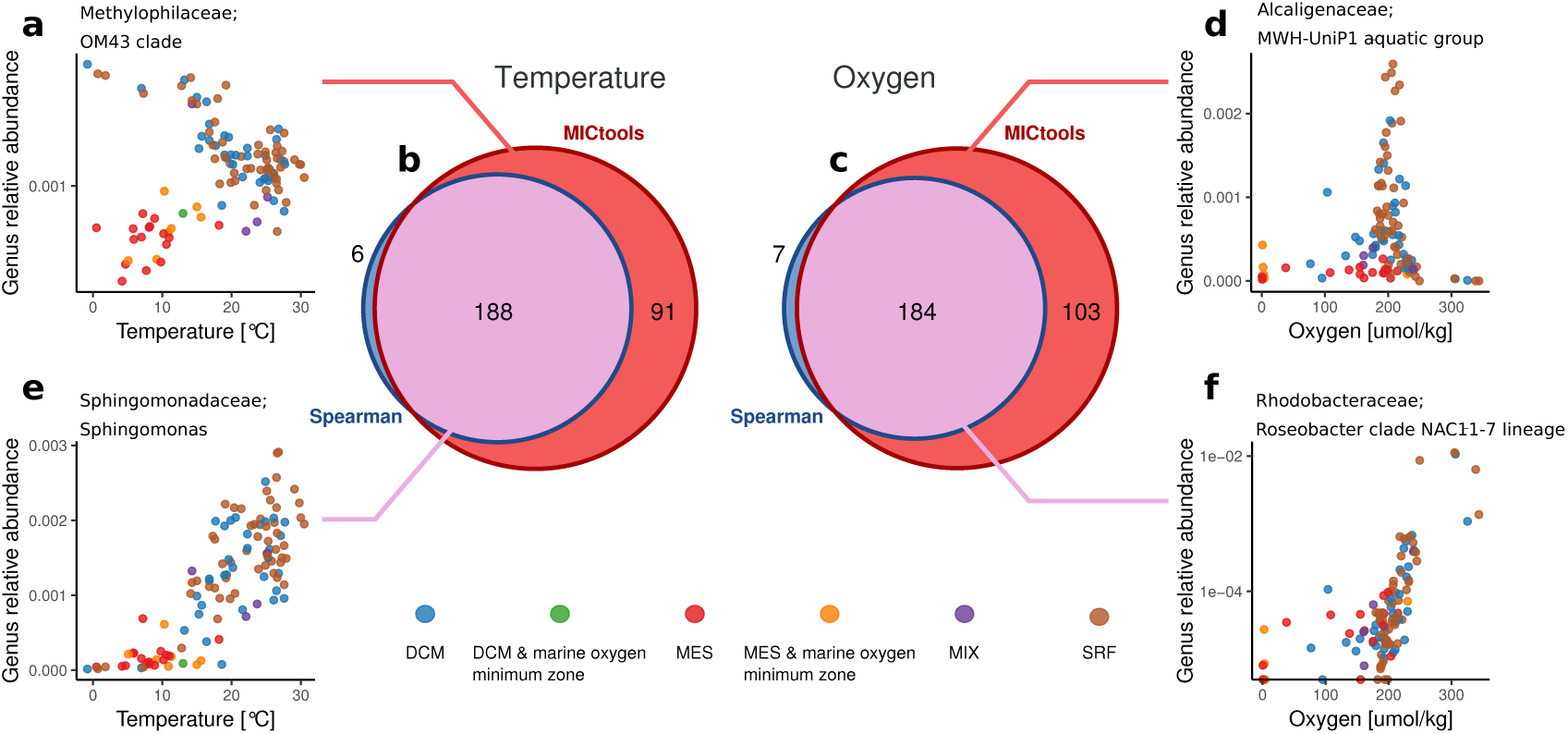
Tara dataset: venn diagrams of the significant relationships between the genus-level relative abundances and two environmental variables, temperature (b) and oxygen (c) identified by MICtools and the Spearman-based procedure (*q* < 0.01). (a, d): the relationships between the OM43 clade and the temperature and between the MWH-UniP1 aquatic group are detected only by MICtools. (e, f): two monotonic relationships identified by both methods. Abbreviations: DCM, deep chlorophyll maximum layer; MES, mesopelagic zone; MIX, subsurface epipelagic mixed layer; SRF, surface water layer.

## Implementation details

MICtools is a Python-based open source software (licensed under GPLv3). MICtools requires the minepy [10] (https://minepy.readthedocs.io), Statsmodels [24] and the NumPy, SciPy, pandas and Matplotlib scientific libraries. MICtools can handle different types of experiments:

- given a single dataset X with variables and samples, MICtools evaluates the 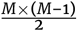 possible associations;
- given two datasets, X (of size *M* × *n*) and Y (of size *K* × *n*) MICtools evaluates all the pairwise relationships between the variables of the two datasets (for a total of *M* × *K* associations).
- given two datasets, X (of size *M* × *n*) and Y (of size *K* × *n*) it evaluates all the row-wise relationships, i.e. only the variables pairs *x*_*i*_ and *y*_*i*_ (for *i* = min(*M, K*)) will be tested;
- moreover, for each experiment listed above, if the sample classes are provided, the analysis will be performed within each class independently.

For multiple testing correction MICtools makes available all the strategies implemented in Statsmodels and a Python implementation of the Storey’s *q*-value method [7]. The indicative number of relationships tested per second during the empirical null estimation (using the TIC_e_) and the strength estimation (MIC_e_) for an increasing number of samples are reported in Additional File 2, Figure A3.

MICtools source and the documentation is available at https://github.com/minepy/mictools. The Docker (https://www.docker.com/) image, containing MICtools and the minepy library is available at https://hub.docker.com/r/minepy/mictools/ and installable with the command `docker pull minepy/mictools`.

## Availability of source code and requirements

- Project name: MICtools
- Project home page: https://github.com/minepy/mictools
- Operating system(s): Platform independent
- Programming language: Python
- Other requirements: minepy, Statsmodels, NumPy, SciPy, pandas, Matplotlib
- License: GNU GPLv3

## Availability of supporting data and materials

The Tara dataset is available at http://ocean-microbiome.embl.de/companion.html.

## Declarations

### NOMENCLATURE

FDR: False Discovery Rate
FN: False Negative
FP: False positive
FWER: Family-Wise Error Rate
MIC: Maximal Information Coefficient
SD1: Synthetic dataset 1
SD2: Synthetic dataset 2
TIC: Total Information Coefficient

## Ethical Approval (optional)

Not applicable.

### Consent for publication

Not applicable.

### Funding

This research was supported by the Autonomous Province of Trento (Accordo di Programma).

### Author’s Contributions

D.A., S.R., C.D. and P.F. conceived the manuscript. D.A. and P.F. developed the methodology. D.A. wrote the software. DA, SR and C.D analyzed the data. D.A., S.R., C.D. and P.F. wrote the manuscript.

## Acknowledgements

Not applicable.

## References

1. Reshef DN, Reshef YA, Finucane HK, Grossman SR, McVean G, Turnbaugh PJ, et al. Detecting Novel Associations in Large Data Sets. Science 2011;334(6062) 1518–1524.

2. Kinney JB, Atwal GS. Equitability, mutual information, and the maximal information coefficient. Proceedings of the National Academy of Sciences 2014;111(9) 3354–3359.

3. Murrell B, Murrell D, Murrell H. R2-equitability is satisfiable. Proceedings of the National Academy of Sciences 2014;111(21):E2160–E2160.

4. Reshef DN, Reshef YA, Mitzenmacher M, Sabeti PC. Cleaning up the record on the maximal information coefficient and equitability. Proceedings of the National Academy of Sciences 2014;111(33):E3362–E3363.

5. Reshef YA, Reshef DN, Finucane HK, Sabeti PC, Mitzenmacher M. Measuring Dependence Powerfully and Equitably. J Mach Learn Res 2016;17(212):1–63.

6. Reshef YA, Reshef DN, Sabeti PC, Mitzenmacher MM. Equitability, interval estimation, and statistical power. arXiv preprint 2015 May;.

7. Storey JD, Tibshirani R. Statistical significance for genomewide studies. Proc Natl Acad Sci U S A 2003 Aug;100(16):9440–9445.

8. Franceschi P, Giordan M, Wehrens R. Multiple comparisons in mass-spectrometry-based-omics technologies. Trends Analyt Chem 2013;50:11–21.

9. Simon N, Tibshirani R. Comment on “Detecting Novel Associations In Large Data Sets” by Reshef Et Al, Science Dec 16, 2011 2014 Jan;.

10. Albanese D, Filosi M, Visintainer R, Riccadonna S, Jurman G, Furlanello C. minerva and minepy: a C engine for the MINE suite and its R, Python and MATLAB wrappers. Bioinformatics 2012;29(3):407–408.

11. Storey JD. A direct approach to false discovery rates. J R Stat Soc Series B Stat Methodol 2002;64(3):479–498.

12. Reshef DN, Reshef YA, Sabeti PC, Mitzenmacher MM. An Empirical Study of Leading Measures of Dependence. arXiv preprint 2015 May;.

13. North BV, Curtis D, Sham PC. A note on the calculation of empirical P values from Monte Carlo procedures. Am J Hum Genet 2002 Aug;71(2):439–441.

14. Krzywinski M, Altman N. Points of significance: Comparing samples—part I. Nat Methods 2014 Mar;11(3):215–216.

15. IG, A E. An Introduction to Feature Extraction. In: IG, M N, S G, LA Z, editors. Feature Extraction. Studies in Fuzziness and Soft Computing, vol. 207 Springer; 2006.

16. Guyon I, Gunn S, Nikravesh M, Zadeh LA. Feature Extraction: Foundations and Applications. Springer; 2008.

17. Bork P, Bowler C, de Vargas C, Gorsky G, Karsenti E, Wincker P. Tara Oceans. Tara Oceans studies plankton at planetary scale. Introduction. Science 2015 May;348(6237):873.

18. Reshef D, Reshef Y, Mitzenmacher M, Sabeti P. Equitability analysis of the maximal information coefficient, with comparisons. arXiv preprint arXiv:13016314 2013;.

19. Pedregosa F, Varoquaux G, Gramfort A, Michel V, Thirion B, Grisel O, et al. Scikit-learn: Machine Learning in Python. Journal of Machine Learning Research 2011;12:2825–2830.

20. Krzywinski M, Birol I, Jones SJM, Marra MA. Hive plots-rational approach to visualizing networks. Brief Bioinform 2012 Sep;13(5):627–644.

21. Logares R, Sunagawa S, Salazar G, Cornejo-Castillo FM, Ferrera I, Sarmento H, et al. Metagenomic 16S rDNA Illumina tags are a powerful alternative to amplicon sequencing to explore diversity and structure of microbial communities. Environ Microbiol 2014 Sep;16(9):2659–2671.

22. Sunagawa S, Coelho LP, Chaffron S, Kultima JR, Labadie K, Salazar G, et al. Ocean plankton. Structure and function of the global ocean microbiome. Science 2015 May;348(6237):1261359.

23. Best DJ, Roberts DE. Algorithm AS 89: The Upper Tail Probabilities of Spearman’s Rho. Appl Stat 1975;24(3):377.

24. Seabold S, Perktold J. Statsmodels: Econometric and statistical modeling with python; 2010.

